# The cerebellum supports two systems for understanding others in early childhood

**DOI:** 10.64898/2026.08.03.742430

**Authors:** Aikaterina Manoli, Nilsu Sağlam, Sofie L. Valk, Charlotte Grosse Wiesmann

**Author notes:** Correspondence: Aikaterina Manoli, Stephanstrasse 1A, 04103 Leipzig, Germany.

## Abstract

The cerebellum has increasingly been implicated in Theory of Mind (ToM), a hallmark of human social cognition. Yet its role in the development of social understanding remains poorly understood, despite evidence linking early cerebellar disruptions to profound social cognitive deficits. Although explicit ToM reasoning emerges only around four years of age, preverbal infants already excel at predicting others’ actions, raising the question of how the cerebellum supports social cognition early in development. Here, we investigated structural cerebellar correlates of ToM (explicit false belief understanding) and nonverbal action prediction in children aged 3–4 years, a critical developmental period during which explicit ToM emerges. Greater gray matter volume in Crus II, a core node of the adult cerebellar ToM network, was associated with better explicit ToM performance. By contrast, nonverbal action prediction was linked to distinct, non-overlapping clusters in inferior lobule VIIB and the posterior vermis, regions implicated in action observation and salience processing in adults. These cerebellar regions further exhibited dissociable patterns of covariance with cerebral networks, linking Crus II to the canonical ToM network and VIIB/vermal regions to salience-related areas. Our findings reveal neuroanatomically distinct cerebellar substrates supporting two separable components of early social cognition: one anchored to explicit mental-state reasoning and another to an earlier-emerging nonverbal action prediction system potentially grounded in the processing of salient social cues. Together, these findings identify the cerebellum as a key contributor to multiple stages of social cognitive development and suggest that distinct cerebellar systems scaffold the emergence of mature social understanding.

## Introduction

Human social interaction depends fundamentally on the ability to infer the mental states of others. This capacity, known as Theory of Mind (ToM), enables the attribution of thoughts, beliefs, and intentions to other people and is a defining feature of human social cognition^1,2^. Converging evidence suggests that the neural basis of ToM extends beyond the cerebral cortex^3–5^, positioning the cerebellum as an important contributor to the development of this ability early in life^6–8^. However, most research has focused on mature, verbally expressed ToM, overlooking evidence that preverbal infants and young children already excel at anticipating others’ actions and thus appear to consider others’ goals and perspectives before they can explicitly reason about mental states^9–12^. It therefore remains unclear whether cerebellar contributions to social cognition are specific to explicit ToM or also extend to earlier-developing forms of social understanding. Addressing this question is particularly important given that cerebellar damage acquired during the first years of life often results in profound and enduring social difficulties that exceed those associated with adult-onset lesions, suggesting that the cerebellum may be crucial for multiple stages of social cognitive development beyond mature mental state reasoning^8,13,14^.

The emergence of ToM marks a major milestone in social cognitive development. Traditionally, explicit ToM has been associated with a developmental breakthrough between 3 and 5 years of age, when children begin to verbally reason about others’ mental states, as demonstrated by successful performance on false belief tasks, widely considered the hallmark test of mature ToM abilities^15–17^. In these tasks, children are asked to predict how a story character holding a false belief about an object’s location, contents, or identity will act toward that object^15,17^. However, a growing body of evidence suggests that action understanding is present much earlier in nonverbal form. In nonverbal paradigms, preverbal infants and young children appear able to track others’ intentions and perspectives, as for example evidenced by their ability to correctly look at the location where they expect an agent to act^9–12^. These findings raise the question of whether nonverbal action prediction reflects an early form of ToM continuous with later explicit mental state reasoning or instead relies on a distinct system that supports social prediction before the emergence of mature, explicit ToM^17–21^.

On a cortical level, explicit ToM and nonverbal action prediction appear to rely on distinct neural substrates. The emergence of explicit ToM between 3 and 5 years of age is associated with the structural and functional maturation of the canonical ToM network, including frontal and temporoparietal regions such as the dorsal and ventral medial prefrontal cortex (d/vmPFC), temporoparietal junction (TPJ), superior temporal sulcus (STS), and precuneus (PreC)^22–24^. Performance on explicit false belief tasks has been linked to developmental increases in cortical thickness, surface area, and white matter integrity within these regions, as well as greater functional specialization and connectivity across the ToM network^25–27^. By contrast, nonverbal action prediction has been associated with increased surface area of the supramarginal gyrus (SMG) and greater white matter connectivity between the SMG and anterior insula (AI)^26,28^. These regions have been implicated in action observation and constitute core components of the salience network, which is involved in the detection of and attentional orienting toward salient stimuli^29^. Together, these findings raise the possibility that early nonverbal action prediction and later explicit ToM rely on partially distinct neurocognitive mechanisms, with the former potentially depending more strongly on processes involved in attending to socially relevant cues and the latter on increasingly abstract representations of others’ mental states^15,28^.

Beyond the cerebral cortex, the cerebellum is increasingly recognized as an important contributor to explicit ToM. In adults, the posterior cerebellum, particularly Crus I–II, is consistently engaged during explicit false belief reasoning, implicating these regions in mature ToM processing^4–6,30,31^. This account is supported by converging evidence demonstrating extensive functional and anatomical connectivity between the posterior cerebellum and the cerebral ToM network. Recent studies have identified bidirectional task-related interactions and structural cerebello-cerebral loops linking posterior cerebellar regions with canonical ToM areas^6,32–34^, suggesting that the cerebellum contributes to explicit mental state reasoning through close interactions with the cerebral ToM network.

The role of the cerebellum in explicit ToM may be particularly important during development. Increasing recruitment of Crus I–II and stronger functional coupling with cerebral ToM regions accompany the emergence of explicit false belief reasoning in early childhood^6^. Moreover, task-based connectivity analyses indicate that cerebellar influences on the cerebral ToM network may be stronger in childhood than in adulthood, raising the possibility that the cerebellum contributes to the establishment of the cognitive processes supporting explicit mental state reasoning^6,8^. Consistent with the proposed role of the cerebellum as a “universal learning system” that constructs and refines internal models of actions and behaviors across cognitive domains^35–37^, these findings suggest that the cerebellum may support explicit ToM by generating predictive representations of others’ mental states, thereby scaffolding the emergence of mature social understanding^8,38,39^.

Despite growing evidence for the cerebellum’s role in the development of explicit ToM, it remains unknown whether cerebellar contributions are specific to mature mental state reasoning or also extend to earlier-developing, nonverbal forms of social understanding. Specifically, it is unclear whether posterior cerebellar regions such as Crus I–II, which are consistently implicated in explicit ToM, also support early nonverbal action prediction, or whether these abilities rely on distinct cerebellar systems, paralleling dissociations observed in the cerebral cortex. Resolving this question would provide important insight into whether early nonverbal action prediction and later explicit mental state reasoning arise from shared or partially distinct developmental mechanisms. This issue is particularly important because converging evidence suggests that the cerebellum plays a distinctive role in social cognitive development more broadly. Cerebellar growth explains unique variance in social behavior beyond that accounted for by the cerebral cortex^7^. Moreover, cerebellar abnormalities, particularly when occurring during the first months of life^8,40^, are consistently associated with persistent social difficulties and neurodevelopmental conditions characterized by atypical social functioning, including autism^14,41,42^. Addressing this question is therefore critical for understanding the developmental foundations of human social cognition and for identifying cerebellar systems that may be relevant to neurodevelopmental disorders involving social difficulties.

In this study, we therefore investigated whether the cerebellum supports multiple stages of social cognitive development by examining its structural correlates of explicit ToM and earlier-developing nonverbal action prediction in preschool-aged children. To this end, we leveraged structural magnetic resonance imaging (MRI) and behavioral data from children aged 3–4 years, spanning the critical developmental period during which explicit ToM abilities emerge^26^. Using voxel-based morphometry (VBM), we examined associations between cerebellar gray matter volume and performance on out-of-scanner explicit false belief and nonverbal action prediction tasks^17,26^. Given extensive cerebello-cerebral interactions in social cognition, we further investigated whether cerebellar structure covaries with cerebral morphology as a function of performance on each task. As ToM development is closely linked to children’s broader cognitive development, including executive function and language, all analyses controlled for general cognitive, executive, and linguistic abilities. To determine whether cerebellar contributions to explicit false belief and nonverbal action prediction were dissociable, we additionally controlled for performance on the alternate task.

## Results

### Cerebellar Crus II volume is associated with emerging ToM abilities

We first sought to identify cerebellar gray matter variation associated with children’s explicit ToM development. To this end, we utilized T1-weighted (T1w) MRI data of 37 3–4-year-old children who performed an out-of-scanner explicit false belief task (henceforth referred to as the ToMAP dataset). VBM analyses were performed across the entire cerebellar cortex using Jacobian-modulated T1w images. A general linear model estimated associations between ToM performance and cerebellar gray matter volume while controlling for children’s biological sex, chronological age, and total cerebellar volume. Higher ToM performance was associated with greater gray matter volume in a cluster located in left Crus II [peak MNI coordinates: −14 −84 −39; *p*_uncorr_ < .001, corrected for the false discovery rate (FDR^43^) at *q* = .05 (henceforth for all clusters); **Fig. 1A**]. This association remained significant when additionally controlling for general cognitive, executive, and linguistic abilities (see **Methods: Behavioral task battery**) and performance on the nonverbal action prediction task, indicating that it was specific to explicit ToM performance and not attributable to development of broader cognitive abilities or shared variance with earlier-developing social cognitive abilities (**Supplementary Fig. 1**).

**Figure 1.**
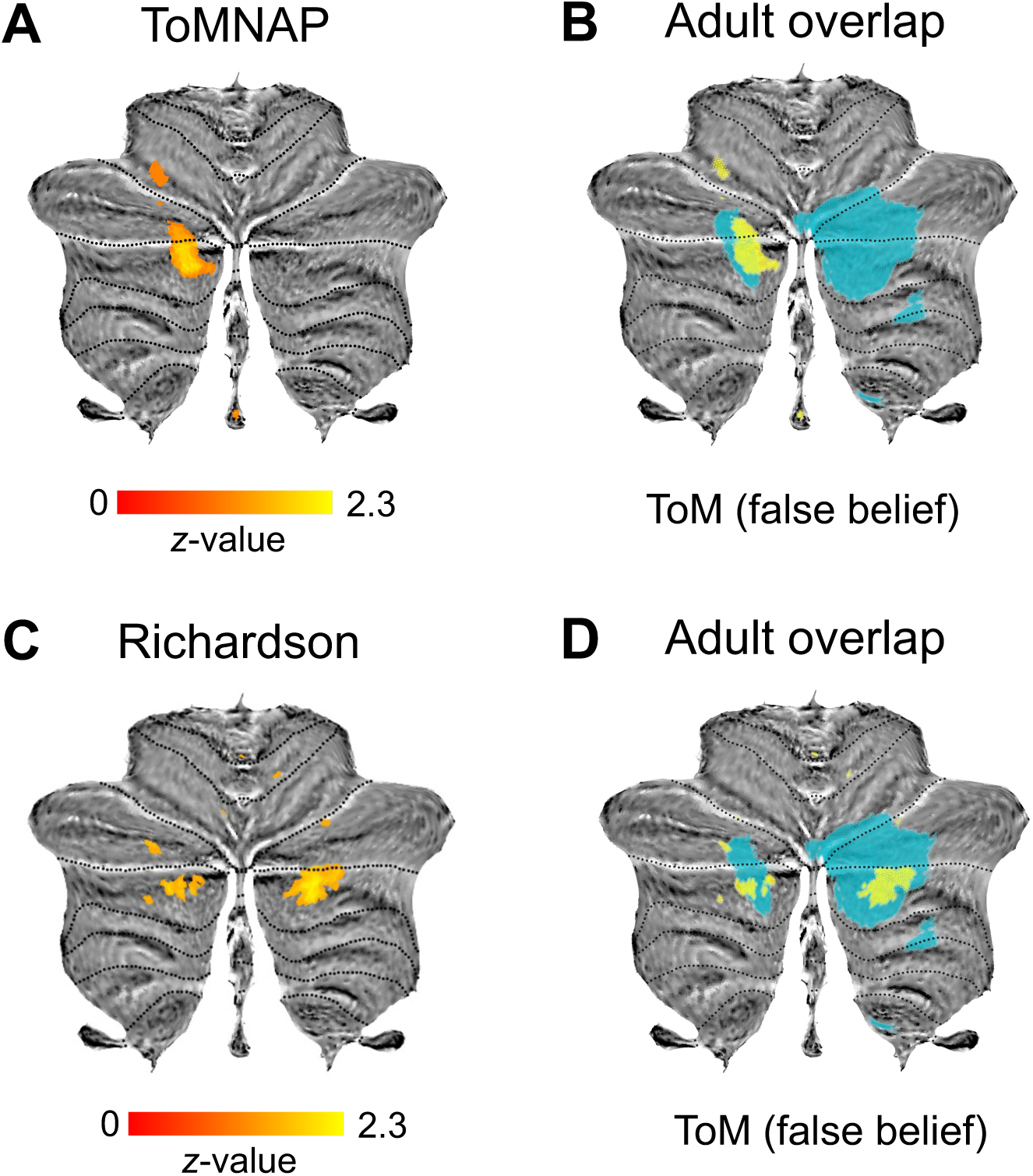
Structural correlates of explicit ToM in the developing cerebellum. **A, C.** Grey matter increases associated with explicit ToM task performance in two independent datasets, ToMAP (**A**) and Richardson (**C**), controlling for age, sex, and total cerebellum size, *z*-scored and FDR-corrected at *q* = .05. **B, D.** Overlap of ToM clusters with adult ToM (false belief) in the multi-domain task battery (MDTB) cerebellar atlas4. Child and adult maps are presented in yellow and blue, respectively.

To assess the robustness of this finding, we replicated the analyses in an independent developmental dataset (Richardson et al.^27^, *N* = 41) employing a similar explicit false belief task, again controlling for children’s sex, age, and total cerebellar volume^6,27^. Consistent with the primary analysis, higher ToM scores were associated with greater gray matter volume in bilateral Crus II (left: −23 −85 −43; right: 24 −80 −39; **Fig. 1C**). In both datasets, the identified Crus II clusters showed substantial spatial overlap with the false belief map derived from the adult multi-domain task battery (MDTB) cerebellar atlas^4^ (**Fig. 1B–D**). This indicates that the current developmental findings converge on a cerebellar territory consistently implicated in explicit ToM processing.

### Lobule VIIB and posterior vermis volume are associated with nonverbal action prediction

A separate general linear model examining nonverbal action prediction performance in the ToMAP dataset, controlling for chronological age, biological sex, and total cerebellar volume, identified significant clusters in bilateral lobule VIIB (left: −13 −75 −52; right: 18 −78 −52) and the posterior vermis (−1 −66 −38; **Fig. 2A**). Notably, these clusters did not overlap with the Crus II regions associated with explicit ToM performance. Given the proposed involvement of salience processing and action observation systems in nonverbal action prediction^26,28^, we compared the identified clusters with the corresponding adult cerebellar maps. The clusters showed spatial correspondence with the inferior cerebellar salience network^44^ (**Fig. 2B**) and action observation map derived from the adult MDTB cerebellar atlas^4^ (**Fig. 2C**). As in the explicit ToM analyses, these associations remained significant when additionally controlling for children’s general cognitive, executive, and linguistic abilities as well as false belief task performance (**Supplementary Fig. 2**), indicating that they were not attributable to broader cognitive abilities or shared variance with explicit ToM.

**Figure 2.**
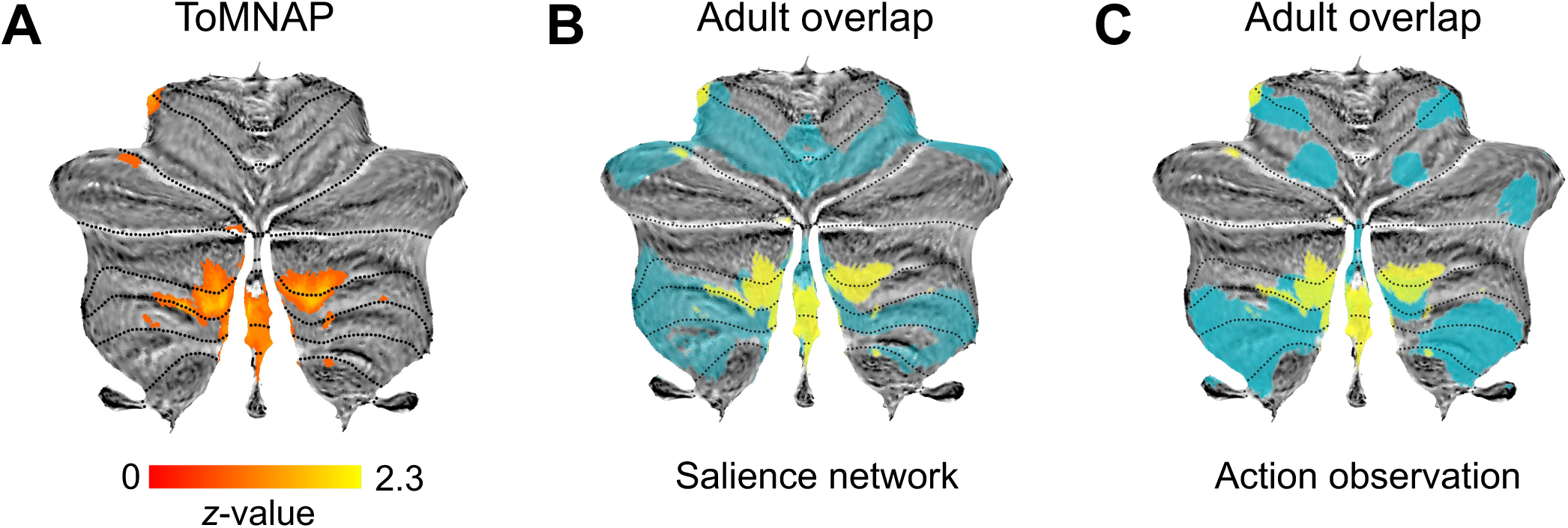
Structural correlates of nonverbal action prediction in the developing cerebellum. **A.** Grey matter increases associated with nonverbal action prediction in the ToMAP dataset, controlling for age, sex, and total cerebellum size, *z*-scored and FDR-corrected at *q* = .05. **B, C.** Overlap of nonverbal action prediction clusters with the adult salience resting-state network44 (**B**) and the action observation map in the multidomain task battery (MDTB) cerebellar atlas4 (**C**). Child and adult maps are presented in yellow and blue, respectively

No other developmental MRI dataset including nonverbal action prediction measures exists to date, so that an independent replication is currently not possible. We therefore assessed the internal robustness of the VBM findings using leave-one-out and bootstrap resampling (**Supplementary Fig. 3**). Across all 37 leave-one-out analyses, every voxel within the original nonverbal action prediction clusters retained a positive association with task performance, indicating that the findings were not driven by any individual participant. Bootstrap analyses (2,000 resamples) further demonstrated high sampling stability, with the association remaining positive in a mean of 99.4% of resamples across the original clusters (median = 99.8%, minimum = 86.7%).

We further investigated cerebellar regions associated with joint attention, another early-developing social ability considered a developmental precursor of later ToM^45,46^, in an independent sample of 1–2-year-old children (Baby Connectome Project; *N* = 31^47^; see **Supplementary Methods**). For joint attention, VBM analyses revealed the largest associations in left lobule VIIB (−26 −73 −50), right lobule VIIIA (8 −70 −53), and right Crus I (44 −63 −39; **Supplementary Fig. 4A**). Interestingly, these regions, particularly lobules VIIB/VIIIA, showed considerable overlap with cerebellar salience and action observation maps identified in adults, mirroring the pattern observed for nonverbal action prediction (**Supplementary Fig. 4B-C**). The findings of inferior posterior cerebellar territories linked to action observation and salience processing both for nonverbal action prediction and joint attention suggest the broader relevance of these regions to early social cognitive abilities.

### Cerebellar ToM and nonverbal action prediction regions show dissociable cerebral covariance profiles

Next, we examined whether the cerebellar regions associated with explicit ToM and nonverbal action prediction exhibited distinct patterns of structural covariance with the cerebral cortex. To this end, we defined 5-mm spherical regions of interest (ROIs) centered on the peak coordinates in left Crus II (explicit ToM) and in bilateral lobule VIIB and the posterior vermis (nonverbal action prediction). General linear models were estimated with voxelwise Jacobian-modulated cerebral gray matter volume as the dependent variable and the interaction between task performance and cerebellar ROI gray matter volume as the predictor of interest, while controlling for biological sex, chronological age, and total intracranial volume. Positive interactions indicate that higher ToM or nonverbal action prediction performance is associated with stronger cerebello-cerebral structural coupling.

For ToM, the interaction between left Crus II gray matter volume and ToM performance revealed significant clusters in the right TPJ and bilateral STS (two-tailed *p*_uncorr_ < .001, FDR-corrected at *q* = .05; **Fig. 3A**, middle). These regions form key components of the canonical ToM network and showed substantial spatial overlap with a recent meta-analysis of ToM in adults^23^ (**Fig. 3A**, right). Additional clusters were observed in the left parietal cortex as well as right-lateralized sensorimotor regions. Importantly, the main findings were replicated in the independent Richardson dataset^27^. Using the same left Crus II seed from the ToMAP dataset, we observed a similar, albeit less extensive, pattern of covariance, including the left STS/superior temporal gyrus (STG), as well as the left PreC of the canonical ToM network (**Fig. 3B**). Peak coordinates and corresponding functional decoding derived using Neurosynth^48^ (https://neurosynth.org/) are reported in **Supplementary Tables 1-2**.

**Figure 3.**
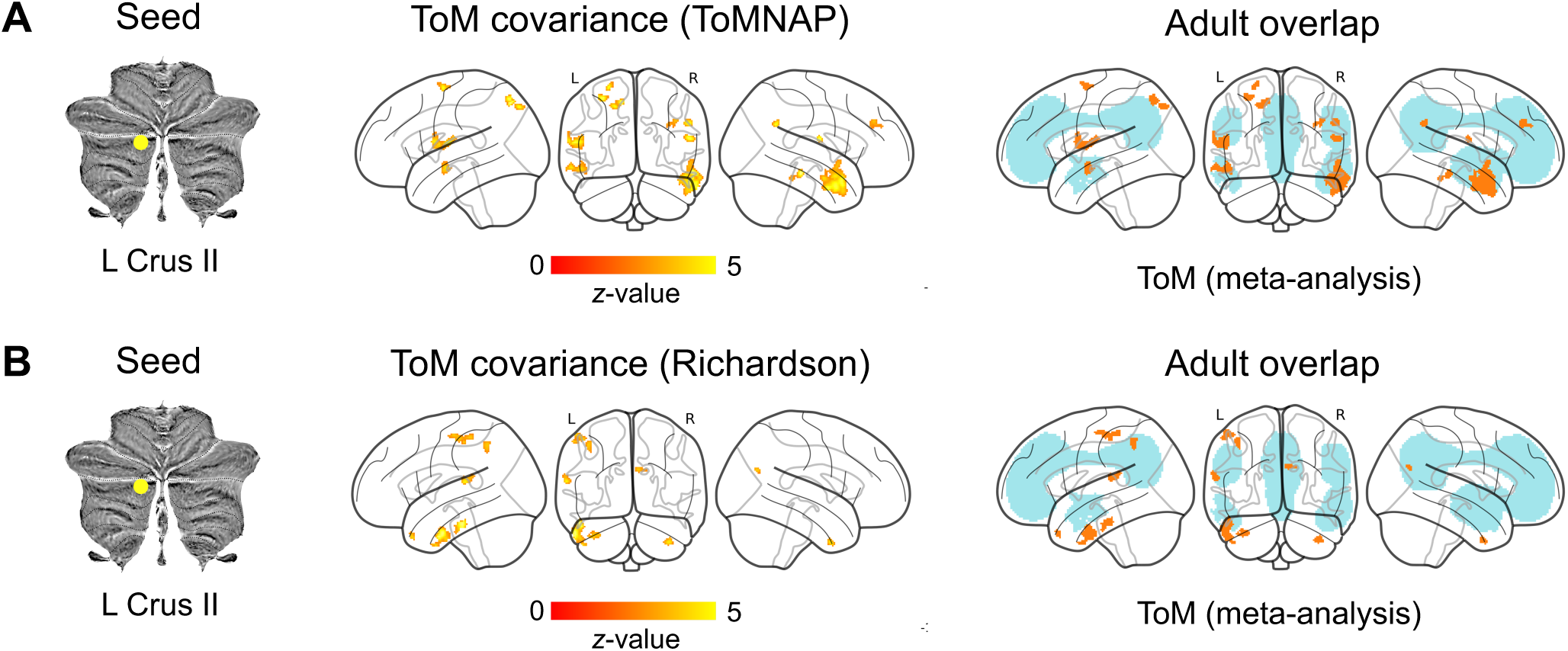
Structural covariance between explicit ToM cerebellar ROI (left Crus II) and cerebral gray matter. **A, B. Left:** ToM cerebellum seed (depicted in yellow). **Middle:** Cerebello-cerebral covariance as a function of ToM in two independent datasets, ToMAP (**A**) and Richardson (**B**), controlling for age, sex, and total cerebellum size, *z*-scored and FDR-corrected at *q* = .05. **Right:** Overlap of explicit ToM covariance clusters with meta-analytic adult ToM23. Child and adult maps are presented in orange and blue, respectively. Abbreviations: L = left.

For nonverbal action prediction, the interaction between each cerebellar ROI and task performance yielded largely consistent cerebello-cerebral covariance clusters in the cingulate cortex, prefrontal cortex, premotor cortex, and somatosensory cortex (two-tailed *p*_uncorr_ < .001, FDR-corrected at *q* = .05; **Fig. 4**, right). Together, these clusters spanned cerebral regions implicated in salience and action perception, memory, and semantic processing. Notably, these clusters did not spatially overlap with the ToM covariance clusters (**Supplementary Fig. 5**). Consistent with previous findings in the cerebral cortex^26,28^ and with the proposed involvement of action observation and salience processing systems in nonverbal action prediction, the bilateral VIIB—and, to a lesser extent, the vermis—covariance clusters overlapped with the cerebral salience network^49^. The left VIIB and, to a greater extent, the vermis additionally covaried with the medial orbitofrontal cortex, a region implicated in nonverbal action prediction, outcome evaluation, and adaptive behavior^50–52^. Interestingly, the vermis also covaried with the STS, a region implicated in biological motion processing, action perception, and ToM^23,53,54^, suggesting that cerebello-cerebral covariance associated with nonverbal action prediction extends beyond salience-related regions to encompass regions broadly involved in the integration and interpretation of socially relevant cues (**Fig. 4**, left). Peak MNI coordinates and corresponding functional associations derived from Neurosynth are reported in **Supplementary Table 3**.

**Figure 4.**
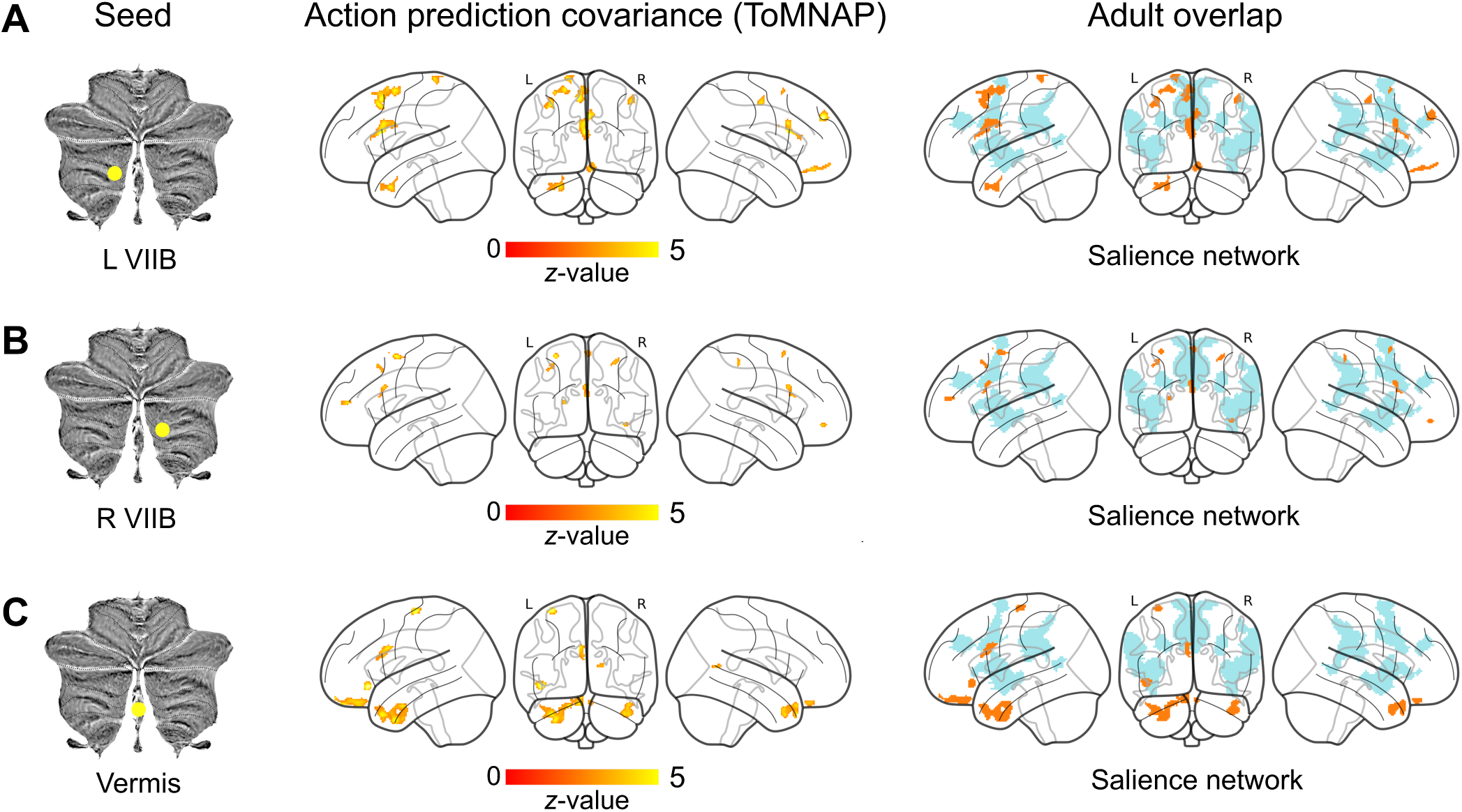
Structural covariance between nonverbal action prediction cerebellar ROIs and cerebral gray matter. **A-C. Left:** nonverbal action prediction cerebellum seeds (depicted in yellow). **Middle:** Cerebellocerebral covariance as a function of nonverbal action prediction in the ToMAP dataset for the left VIIB (**A**), right VIIB (**B**), and vermis (**C**) seeds, controlling for age, sex, and total cerebellum size, *z*-scored andFDR-corrected at *q* = .05. **Right:** Overlap of nonverbal action prediction covariance clusters with the adult salience resting-state network49 (**A-C**). Child and adult maps are presented in orange and blue, respectively. Abbreviations: L = left; R = right.

Importantly, the explicit ToM and nonverbal action prediction covariance patterns remained consistent in additional models controlling for children’s linguistic, executive, and general cognitive abilities, and performance on the alternate social cognitive task in the ToMAP dataset (**Supplementary Figs. 6-7**). As in the VBM analyses, overlap with adult maps was assessed post hoc and aimed to contextualize the findings within established cerebral functional territories.

## Discussion

While the role of the cerebellum in ToM is increasingly recognized, its contribution to earlier-developing social cognitive abilities has remained unclear. Here, we addressed this question by examining cerebellar correlates of explicit ToM and nonverbal action prediction in young children in the critical period before and after the emergence of explicit ToM abilities. We identified neuroanatomically dissociable cerebellar substrates associated with these abilities. Explicit ToM was linked to greater gray matter volume in left Crus II, a region overlapping with canonical cerebellar ToM territories in adults^4^, whereas nonverbal action prediction was associated with non-overlapping clusters in bilateral lobule VIIB and the posterior vermis, regions corresponding to adult action observation and salience processing networks^4,44^. These cerebellar loci additionally exhibited distinct patterns of cerebello-cerebral structural covariance: Crus II covaried with core cerebral ToM regions, including the TPJ, PreC, and STS, whereas nonverbal action prediction regions covaried with cerebral territories implicated in salience processing, action representation, and socio-affective functions. Importantly, these associations remained independent of children’s age, sex, brain size, and general cognitive, executive, and linguistic abilities, suggesting that they reflect specific contributions of the cerebellum to different components of early social cognition.

The present findings extend growing evidence implicating the cerebellum in social cognitive development. Previous work has identified the cerebellum as an important contributor to both typical and atypical social functioning^6–8,13,14,40^, with recent evidence suggesting that cerebellar maturation explains unique variance in social behavior beyond that accounted for by the cerebral cortex alone^7^. Here, we show that cerebellar structure is associated not only with the emergence of explicit ToM, extending previous functional findings^6^, but also with earlier-emerging social abilities. Together, these findings suggest that cerebellar contributions to social cognition span multiple stages of development, from early action-based social understanding to later explicit reasoning about others’ mental states, positioning the cerebellum as a key contributor to the maturation of human social cognition. Given the prominent involvement of the cerebellum in conditions characterized by atypical social cognition, such as autism^8,13,14,40^, the present findings may also help refine neurodevelopmental models of atypical social development by identifying distinct cerebellar pathways relevant to these conditions.

The dissociable patterns observed for explicit ToM and non-verbal action prediction mirror previously reported dissociations in the cerebral cortex and further support the existence of separable, distributed neural systems underlying different forms of social understanding in early life^26,28^. Across two independent samples, explicit ToM was associated with Crus II, a region consistently implicated in mental state reasoning in both adults and children^4,6,30,55^, and exhibited structural covariance with canonical cerebral ToM regions^22,23^. Notably, both cerebellar and cerebral ToM regions overlap substantially with the default mode network^44,49,56^, a system implicated in internally generated cognition, episodic simulation, and social reasoning^57–59^. Because these functions depend on representations that are abstract and decoupled from immediate perceptual input^59^, the coordinated maturation of Crus II and cerebral ToM regions may reflect children’s developing capacity to generate and manipulate abstract representations of others’ mental states.

By contrast, nonverbal action prediction was associated with inferior posterior cerebellar regions, particularly lobule VIIB and the posterior vermis. These regions overlapped with adult maps of action observation and salience processing^4,44^ and covaried with cerebral regions implicated in salience detection, action perception, and socio-affective processing (https://neurosynth.org/^48^). Many of these functions—including action understanding, empathy, and understanding another’s visual perspective—emerge earlier in development than explicit ToM reasoning^10,60–62^. One possibility is that successful performance in these tasks relies, at least in part, on heightened sensitivity to “bottom-up” socially informative cues, such as another agent’s actions and gaze direction, rather than on abstract, “top-down” mental state representations^28^. Such mechanisms may enable young children to predict and interpret others’ behavior before the emergence of mature mental state reasoning.

The observed patterns of cerebello-cerebral covariance further suggest that cerebellar contributions to social cognition are embedded within distributed developmental systems linking the cerebellum and cerebral cortex. Structural covariance is thought to reflect coordinated maturation among anatomically and functionally connected brain regions^63,64^. Consistent with this view, previous research has shown that cerebellar morphology closely mirrors that of the cerebral cortex^65^ and that cerebellar and cerebral regions supporting similar cognitive functions co-mature in a domain-specific manner^7^. These findings are particularly relevant in light of developmental accounts proposing that the cerebellum plays a disproportionately important role early in life by shaping the maturation of connected cerebral regions^14^. Accordingly, the dissociable patterns of cerebello-cerebral covariance observed here suggest that the cerebellum may contribute to the development of multiple, partially independent social cognitive systems, with distinct cerebellar territories coordinating the maturation of cerebral networks supporting explicit mental state reasoning and earlier-emerging social prediction abilities.

However, the mechanisms through which the cerebellum contributes to explicit ToM and nonverbal action prediction remain an open question. One influential account proposes that the cerebellum contributes to mature ToM by generating and refining internal models of social interactions, thereby enabling predictions about others’ mental states^35,38,39^. This framework suggests a particularly important role for cerebellar computations during early childhood, when the cerebellum may contribute to the initial acquisition and refinement of predictive models that support explicit mental state reasoning^6,8^. By contrast, the distinct cerebellar correlates of nonverbal action prediction observed here suggest that earlier social abilities may rely on different cerebellar computations, potentially involving the processing of context-sensitive and socially salient perceptual information rather than structured internal representations of others’ mental states^28,66,67^. Future longitudinal, behavioral, and multimodal neuroimaging studies will be necessary to determine how these distinct cerebellar systems contribute to the development of different forms of social understanding.

Additionally, no other independent developmental dataset of nonverbal action prediction was available. However, the present findings proved highly robust to leave-one-out and bootstrap resampling, indicating that they were neither driven by individual participants nor unduly sensitive to sampling variability. We further observed partly overlapping associations in an independent dataset of joint attention, another early-developing social cognitive ability and precursor of explicit ToM^45,46^. Specifically, joint attention was associated with lobule VIIB and overlapped with adult cerebellar salience and action observation maps^4,44^. While these findings do not constitute a direct replication of the present results, they raise the possibility that inferior posterior cerebellar regions may support a broader system underlying early social cognitive abilities, extending beyond nonverbal action prediction alone. Future studies should determine whether these regions represent a common cerebellar substrate for multiple precursors of mature social understanding and, importantly, test the reproducibility of the present findings in independent developmental neuroimaging datasets that include nonverbal action prediction measures.

Lastly, unlike the explicit ToM covariance analysis, where we validated our findings using an independently defined cerebellar seed in the Richardson dataset, no comparable independent seed was available for nonverbal action prediction. Consequently, the structural covariance patterns associated with lobule VIIB and the posterior vermis should be considered hypothesis-generating rather than definitive until replicated using independently defined regions of interest. Relatedly, whereas Crus II has been consistently implicated in the broader cerebral ToM network through functional and anatomical connectivity studies^6,32,33,68^, considerably less is known about how lobule VIIB and the posterior vermis are embedded within broader cerebello-cerebral networks. In particular, the connectivity of these regions with the SMG and other regions of the salience and action observation networks implicated in nonverbal action prediction remains to be established^26,28^. Integrating the present structural covariance findings with functional and anatomical connectivity measures may provide a more comprehensive characterization of the network architecture supporting early forms of social understanding.

Overall, the present study demonstrates dissociable cerebellar correlates of explicit ToM and earlier-developing nonverbal action prediction. Whereas explicit ToM was associated with Crus II, a core node of the cerebellar ToM network, nonverbal action prediction was more strongly linked to inferior cerebellar and vermal regions implicated in action observation and salience processing, as further supported by their covariance with functionally related cerebral territories. These findings highlight the cerebellum as an important contributor to multiple stages of social-cognitive development and support the view that distinct cerebellar systems underpin different forms of social understanding early in life. By extending cerebellar accounts of social cognition beyond explicit mental state reasoning, the present work provides new insight into the neural foundations of social development and may ultimately inform efforts to understand neurodevelopmental conditions characterized by atypical social cognitive processing.

## Methods

### Preregistration

Our research questions and analysis approach were preregistered prior to data analysis (https://aspredicted.org/638fh2.pdf).

### Participants

We utilized T1w MRI and behavioral data from 38 typically developing children which examined the development of ToM and nonverbal action prediction abilities. The in-house ToMAP dataset includes T1w MRI and behavioral data from 3-to 4-year-old children who completed out-of-scanner assessments of ToM (an explicit false belief task) and nonverbal action prediction (a nonverbal anticipatory looking task; see **Behavioral task battery**). One child was excluded from the present analyses due to incomplete behavioral assessments. The final sample consisted of 37 children (age range: 3-4 years; *M* = 3.60 years, *SD* = 0.50; 17 female), all of whom had complete cerebellar coverage. A power analysis with G*Power^69^ showed that the computed linear regression with *N* = 37, an *α* error of 5%, and an effect size of *f* = 0.25^16,26^ had a power of 1-*β* = 84%. Parental informed consent was obtained for all participants, and data collection was approved by the Ethics Committee at the Faculty of Medicine of the University of Leipzig. The present dataset was previously used to address different research questions^25,26,28^. The question of cerebellar involvement in the development of social cognitive abilities has not been addressed before.

The Richardson et al.^27^ dataset, used for replication of the explicit ToM findings, includes T1w, functional, and behavioral data of 122 children who performed an out-of-scanner assessment of ToM abilities, namely a battery of explicit false belief tasks. Children were excluded from the original study if they did not complete the behavioral and neuroimaging components, if they moved excessively in the scanner or if they demonstrated language delays. As in our previous work investigating functional involvement of the cerebellum in ToM development with the same dataset^6^, we further excluded children with insufficient cerebellar coverage in T1w scans after visual inspection (*N* = 71), with our final sample consisting of 41 children (age range: 3-12 years; age: *M* = 5.91 years, *SD* = 2.29; 25 female). Parental informed consent was obtained for all participants in the original study, and data collection was approved by the relevant ethics committee.

### Behavioral task battery

#### Explicit ToM

Explicit ToM abilities were assessed using an out-of-scanner false belief task^25,27^. False belief understanding is widely considered the critical marker of ToM development^15^. In both the ToMAP and Richardson datasets, children completed six canonical false belief questions involving unexpected changes in an object’s location or content while a story protagonist who had previously interacted with the object left the scene^70^. For example, children were asked where a protagonist holding a false belief about an object’s location would search for it upon re-entering the scene, and why. In the ToMAP dataset, false-belief scenarios were interactively enacted in front of the children using animal puppets, whereas in the Richardson dataset, false belief scenarios were presented as verbally narrated stories. Children’s responses were aggregated into a composite ToM score ranging from 0 to 6 correct answers. In the primary ToMAP dataset, performance differed significantly between 3-and 4-year-olds (Mann–Whitney *U* test, *p* < .001, two-tailed). Three-year-olds performed below chance (median = 0; Wilcoxon signed-rank test, *p* < .001), whereas four-year-olds performed marginally above chance (median = 0.50; Wilcoxon signed-rank test, *p* = .082, two-tailed).

#### Nonverbal action prediction

Nonverbal action prediction was assessed using an out-of-scanner anticipatory-looking task from the ToMAP dataset^17^. Children viewed short videos while their gaze was recorded using a Tobii T120 eye-tracker. The videos depicted an animal agent observing a mouse run through a Y-shaped tunnel toward one of two boxes located at the tunnel exits. Following familiarization trials, in which the agent successfully tracked the mouse, children completed false belief trials in which the agent held an incorrect belief about the mouse’s location. Specifically, after observing the mouse enter one of the boxes, the agent left the scene. During the agent’s absence, the mouse exited the scene entirely, leaving the agent with a false belief about its whereabouts. Children’s anticipatory gaze while the agent traversed the tunnel was used to index their prediction of where the agent would search for the mouse. The original task comprised two conditions. In the first condition (FB1), the agent observed the mouse move from one box to the other before leaving the scene and therefore did not witness the mouse subsequently exit the scene.

In the second condition (FB2), the agent left immediately after observing the mouse enter a box and similarly did not observe the mouse leaving the scene (see **Supplementary Fig. 8** for a schematic depiction of the task). In the present study, we focused on condition FB1 because it has shown greater robustness and has been independently replicated in a behavioral sample^21^. However, analyses using a combined FB1–FB2 score originally used in Grosse Wiesmann et al.^26^ yielded highly similar results and are reported in **Supplementary Fig. 9**.

Gaze data were analyzed within a predefined anticipatory window extending from the moment the agent entered the tunnel until the end of the trial. Two areas of interest (AOIs), corresponding to the two tunnel exits, were defined, and looking times toward each exit were quantified. Following established procedures^17,25,71^, a differential looking score reflecting relative attention toward the belief-consistent versus belief-inconsistent location was calculated and used as the action-prediction measure. Across the sample, children looked significantly more to the correct than the incorrect location [*M* = 55.64%, *SD* = 12.11%; one-sample t-test: *t* (36) = 2.83, *p* = .007, two-tailed]. There was no significant age-related difference between 3-and 4-year-olds in how often children looked at the correct versus incorrect location [3-year-olds: *M* = 57.74%, *SD* = 14.95%; 4-year-olds: *M* = 54.05%, *SD* = 9.49%; Welch’s *t* (23.94) = 0.86, *p* = .396, two-tailed]. Furthermore, explicit ToM and nonverbal action prediction were not significantly correlated (Spearman’s rank correlation, *ρ* = −.21, *p* = .221, two-tailed).

#### Additional cognitive assessments

Participants in the primary ToMAP dataset completed additional assessments of cognitive abilities to ensure that any observed effects were specific to ToM rather than reflecting broader cognitive development^72,73^. Specifically, children were administered a standardized measure of language development [*Sprachentwicklungstest Kinder* (SETK)^74^], and a battery of executive function tasks targeting inhibitory control, response selection, and cognitive flexibility, previously shown to be associated with ToM performance^17^. Additionally, children completed an assessment of general cognitive abilities, including memory, attention, and spatial representation^75^. All children completed the full set of assessments, including both ToM and nonverbal action prediction tasks. The order of tasks was counterbalanced across participants, with the exception of the explicit ToM tasks, which were always administered last to prevent potential carryover effects on the nonverbal action prediction task. Detailed descriptions of the tasks can be found in Grosse Wiesmann et al.^17^.

### MRI data acquisition

In the ToMAP dataset, structural MRI data were acquired on a 3T Siemens TIM Trio scanner equipped with a 32-channel head coil. High-resolution whole-brain T1w images were obtained using an MP2RAGE sequence with a voxel size of 1.2 × 1.0 × 1.0 mm (TI1 = 700 ms, flip angle1 = 4°; TI2 = 2,500 ms, flip angle2 = 5°; TR = 5,000 ms; TE = 3.24 ms; field of view = 192 × 192 mm; 176 sagittal slices; GRAPPA acceleration factor = 3; partial Fourier factor = 6/8; bandwidth = 240 Hz/Px; acquisition time = 5 min 22 s). To facilitate successful scanning, children participated in a mock-scanning session several days before data acquisition and watched a self-selected movie using MR-compatible goggles and headphones during the scan.

MRI acquisition procedures for the Richardson dataset have been described previously ^27^. Briefly, whole-brain T1w images were collected on a 3T Siemens Tim Trio scanner. Children younger than five years were scanned using a custom 32-channel phased-array head coil adapted for smaller head sizes, whereas older children were scanned with a standard Siemens 32-channel head coil. Images were acquired in 176 interleaved sagittal slices with 1-mm isotropic resolution using GRAPPA parallel imaging (acceleration factor = 3). The field of view was 192 mm for the child coil and 256 mm for the adult coil. As in the ToMAP dataset, children completed a mock-scanning session before MRI acquisition to familiarize themselves with the scanning environment.

### MRI data analysis

#### Cerebellar segmentation

Across datasets, the cerebellum was isolated and normalized to the Spatially Unbiased Infra-Tentorial (SUIT^76^) template using the SUIT toolbox (v.3.5) (https://github.com/jdiedrichsen/suit/releases/tag/3.5), implemented in SPM12^77^ (http://www.fil.ion.ucl.ac.uk/spm/software/spm12) within MATLAB R2025b (The MathWorks, 2025). Normalization to a spatially unbiased, cerebellum-specific template has been shown to markedly improve alignment of individual cerebellar fissures and deep cerebellar nuclei compared with normalization to standard ICBM152 MNI space (MNI152NLin2009cAsym^78^)^31,76,79,80^. For each participant, the cerebellum was first extracted and segmented into gray and white matter, and a cerebellar isolation mask was generated. These masks were visually inspected and, when necessary, manually corrected for each subject using ITK-SNAP^81^ (v.4.4.0) (https://itksnap.org/pmwiki/pmwiki.php), ensuring that voxels from the inferior temporal and occipital cerebral cortex were excluded. The individual gray and white matter probabilistic maps were then normalized to SUIT space using diffeomorphic anatomical registration through the exponentiated Lie algebra algorithm^82^. This approach deforms the cerebellum to simultaneously align cerebellar gray and white matter probability maps with the SUIT atlas template. The resulting nonlinear deformation was applied to each participant’s cerebellar gray matter maps using the hand-corrected cerebellar masks, with modulation by the Jacobian determinants to preserve regional gray matter volume following normalization^83^. For VBM analyses, the normalized, modulated images were smoothed using a 5-mm FWHM Gaussian kernel. All data were visualized on the cerebellar flatmap, a surface-based reconstruction of the cerebellar cortex that allows the full spatial extent of cerebellar activations to be displayed^31^. As the flatmap does not provide an anatomically accurate representation of extensive, cerebellar folding, it was used exclusively for visualization purposes^4,31^.

#### Cerebral cortical segmentation

For segmentation of the cerebral cortex, T1w images were processed using the Computational Anatomy Toolbox (CAT12^84^; https://neuro-jena.github.io/cat/), implemented in SPM12^77^ (http://www.fil.ion.ucl.ac.uk/spm/software/spm12) within MATLAB R2025b (The MathWorks, 2025). Cerebral segmentation and spatial normalization were performed using the standard CAT12 *estwrite* pipeline, which consists of tissue segmentation followed by spatial registration. Tissue segmentation was initiated by applying a spatially adaptive nonlocal means (SANLM) denoising filter^85^, followed by SPM’s unified segmentation procedure^86^ to obtain initial estimates of gray matter, white matter, and cerebrospinal fluid. These outputs were subsequently refined using CAT12-specific optimization steps. First, the brain was parcellated into the left and right cerebral hemispheres, subcortical structures, ventricles, and cerebellum, and local white matter hyperintensities were detected to account for their influence during later processing. Second, a local intensity transformation was applied to reduce regional intensity inhomogeneities associated with variable degrees of cortical myelination, particularly in the motor cortex, basal ganglia, and occipital cortex. Finally, tissue classification was refined using CAT12’s adaptive maximum a posteriori (AMAP) approach^87^, which does not require a priori tissue probability information and incorporates partial volume estimation to improve tissue boundary definition. Segmented tissue classes were spatially normalized to standard ICBM152 MNI space (MNI152NLin2009cAsym^78^) using CAT12’s nonlinear registration framework based on geodesic shooting^88^, which combines affine and high-dimensional deformation to account for individual anatomical variability. Modulated, normalized tissue probability maps were generated, such that voxel intensities reflect regional tissue volume following normalization. For VBM analyses, the modulated gray matter maps were smoothed using a 5-mm FWHM Gaussian kernel. All CAT12 processing was executed using default parameters, including the standard SPM tissue probability maps, a bias regularization strength of 0.5, and a sampling distance of 3 mm. All segmentations were visually inspected using CAT12’s quality control reports, indicating good to excellent segmentation quality.

#### Voxel-based morphometry

VBM analyses were conducted within modulated cerebellar gray matter maps using the Nilearn library (version 0.10.1; https://nilearn.github.io/stable/index.html) in Python (version 3.9.7) to examine regional associations between cerebellar structure and performance on ToM and nonverbal action prediction tasks. Voxel-wise general linear models were specified with the task score of interest (ToM or nonverbal action prediction) entered as the predictor variable. Children’s biological sex, chronological age, and total cerebellar volume (computed as the sum of normalized cerebellar gray and white matter volumes) were included as additional covariates. To ensure that observed effects were specific to each social cognitive ability, children’s executive function, language, and general cognitive abilities were also included as covariates. To assess the independence of effects between tasks, performance on the ToM task was included as a predictor in the nonverbal action prediction model, and conversely, nonverbal action prediction performance was included as a predictor in the ToM model. All continuous predictors were mean-centered prior to model estimation, and all statistical tests were two-tailed. Statistical significance was determined using voxel-wise FDR correction^43^ at *q* = .05 across the cerebellum.

### Robustness analyses of nonverbal action prediction VBM

Because no independent developmental neuroimaging dataset including nonverbal action prediction measures was available, we conducted additional robustness analyses for the primary VBM findings related to nonverbal action prediction. First, we performed leave-one-out resampling by repeating the complete voxel-wise VBM analysis 37 times, each time excluding one participant. Second, we performed a nonparametric bootstrap analysis with 2,000 resamples, sampling participants with replacement while preserving the original sample size. For both analyses, we calculated voxel-wise positive-frequency maps indicating the proportion of resampled analyses in which the association with nonverbal action prediction remained positive within the clusters identified in the primary analysis. These analyses were designed to assess whether the reported associations were robust to the exclusion of individual participants and to sampling variability in the absence of an independent replication dataset.

### Cerebello-cerebral covariance

Spherical ROIs (5-mm radius) were defined within cerebellar gray matter based on the local maxima identified in the cerebellar VBM analyses. For ToM, an ROI was centered on the left Crus II peak (−14 −84 −39), which was identified in the primary ToMAP dataset and showed substantial overlap with the corresponding cluster in the Richardson dataset. For nonverbal action prediction, ROIs were centered on the bilateral lobule VIIB peaks (left: −13 −75 −52; right: 18 −78 −52) and the posterior vermis peak (−1 −66 −38). These ROIs were applied to the modulated cerebellar gray matter maps and mean gray matter volume within each sphere was extracted for each participant.

To examine cerebello–cerebral structural associations as a function of ToM or nonverbal action prediction score, voxel-wise general linear models were estimated in which cerebral cortical gray matter volume served as the dependent variable. Analyses were conducted using modulated cortical gray matter maps derived from CAT12 processing and were restricted to voxels within an explicit cerebral gray matter mask generated from the mean normalized gray matter image across all participants. The primary effect of interest was the interaction between ToM or nonverbal action prediction task performance and cerebellar ROI gray matter volume, allowing assessment of whether the association between cerebellar structure and cerebral gray matter varied as a function of task performance. A positive interaction indicates that higher task performance is associated with a stronger positive relationship between cerebellar ROI gray matter volume and cerebral gray matter volume, suggesting increased cerebello–cerebral structural coupling in individuals with more advanced ToM or nonverbal action prediction abilities.

All models included children’s biological sex, chronological age, executive function, language abilities, general cognitive abilities, performance on the alternate task, and total intracranial volume as additional covariates. Total intracranial volume was computed as the sum of gray matter, white matter, and cerebrospinal fluid volumes obtained from CAT12 outputs. As in the VBM models, continuous predictors were mean-centered prior to model estimation, and all statistical tests were two-tailed. Statistical significance was assessed using voxel-wise FDR correction^43^ at *q* = .05 across the cerebral cortex.

## Data availability

All materials and data from the ToMAP dataset are stored in a local repository at the Max Planck Institute for Human Cognitive and Brain Sciences. Fully anonymized data are available upon reasonable request, subject to the data protection regulations and ethical approvals governing the study. The ToM replication dataset^27^ is publicly available through OpenNeuro (https://openneuro.org/datasets/ds000228/versions/1.1.0). The joint attention dataset from the Lifespan Baby Connectome Project^47^ is available through the National Institute of Mental Health Data Archive (https://nda.nih.gov/edit_collection.html?id=2848).

Adult maps used to contextualize the developmental findings are publicly available: MDTB cerebellar atlas^4^ (https://github.com/DiedrichsenLab/cerebellar_atlases/tree/master/King_2019), cerebellar resting-state functional networks^44^ (https://github.com/DiedrichsenLab/cerebellar_atlases/tree/master/Buckner_2011), ToM meta-analysis maps^23^ (https://osf.io/pav27/files/mrb35), and cerebral resting-state functional networks^49^ (https://github.com/ThomasYeoLab/CBIG/tree/master/stable_projects/brain_parcellation/Yeo20 11_fcMRI_clustering/1000subjects_reference/Yeo_JNeurophysiol11_SplitLabels).

## Code availability

Data processing and segmentation leveraged openly available software: SUIT for cerebellar segmentation (https://github.com/jdiedrichsen/suit/releases/tag/3.5) and CAT12 for cerebral segmentation (https://neuro-jena.github.io/cat/), both implemented in SPM12 (http://www.fil.ion.ucl.ac.uk/spm/software/spm12). All scripts used for image processing, statistical analyses, and figure generation are publicly available on GitHub (https://github.com/kmanoli/cereb_tom_anat).

## Supporting information

Supplementary information for manuscript.

## Acknowledgments

We thank Jörn Diedrichsen, Arno Villringer, and Jessica Royer for insightful comments on data analysis and results interpretation. This work is supported by the Max Planck Society. AM was also funded by the German Academic Scholarship Foundation (*Studienstiftung des deutschen Volkes*) and the Lise Meitner Excellence Program (awarded to SLV). NS was supported by Erasmus+ and the Lise Meitner Excellence Program (awarded to SLV). SLV was also funded in part by Helmholtz Association’s Initiative and Networking Fund under the Helmholtz International Lab grant agreement InterLabs-0015, and the Canada First Research Excellence Fund (CFREF Competition 2, 2015-2016) awarded to the Healthy Brains, Healthy Lives initiative at McGill University, through the Helmholtz International BigBrain Analytics and Learning Laboratory (HIBALL). SLV was furthermore supported by the Jacobs Foundation Research Fellowship, the Hector Research Career Development Award, and a European Research Council (ERC) Starting Grant (SOCO). CGW was supported by an ERC Starting Grant (REPRESENT 101117806).

## Author contributions

AM, SLV and CGW conceptualized and designed the study. AM and NS processed and analyzed the data with input from SLV and CGW. AM created the visualizations and wrote the original draft. AM, NS, SLV, and CGW contributed to the interpretation of results. AM, NS, SLV, and CGW edited and reviewed the final manuscript. SLV and CGW contributed equally and share senior authorship.

