## Supplementary information for manuscript. for "The cerebellum supports two systems for understanding others in early childhood"

* = Shared last authorship.

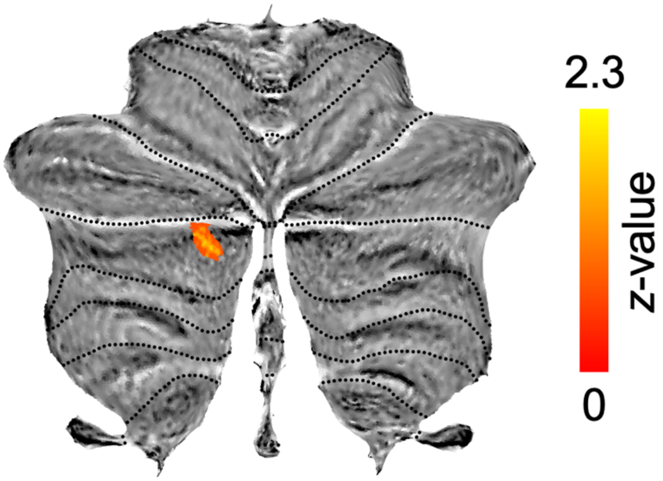


**Supplementary Figure 1.** Grey matter increases associated with explicit ToM task performance, controlling for age, sex, total cerebellum size, general cognitive, executive, and linguistic abilities, and nonverbal action prediction task performance (*z*-scored and FDR-corrected at $q = .05$).

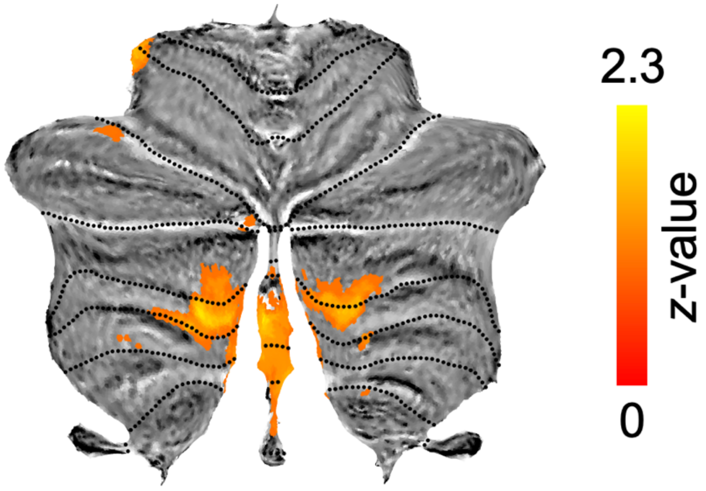


**Supplementary Figure 2.** Grey matter increases associated with nonverbal action prediction task performance, controlling for age, sex, total cerebellum size, general cognitive, executive, and linguistic abilities, and explicit ToM task performance (*z*-scored and FDR-corrected at $q = .05$).

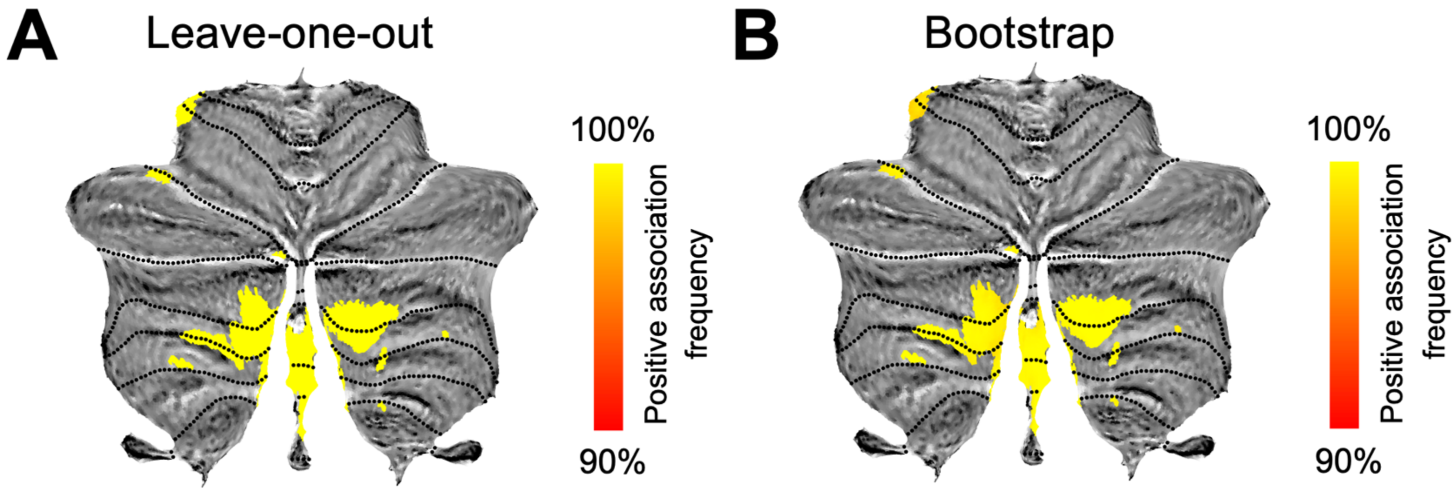


**Supplementary Figure 3.** Internal robustness of cerebellar VBM findings associated with nonverbal action prediction. (**A**) Leave-one-out analysis, in which the complete voxel-wise VBM model was repeated 37 times, each time excluding one participant. (**B**) Nonparametric bootstrap analysis based on 2,000 resamples. Colors indicate the proportion of resampled analyses in which each voxel within the original nonverbal action prediction clusters retained a positive association with task performance. Across all leave-one-out analyses, every voxel within the original clusters retained a positive association (100% directional consistency). Bootstrap analyses likewise demonstrated high sampling stability, with the association remaining positive in a mean of 99.4% of resamples across the original clusters (median = 99.8%, minimum = 86.7%).

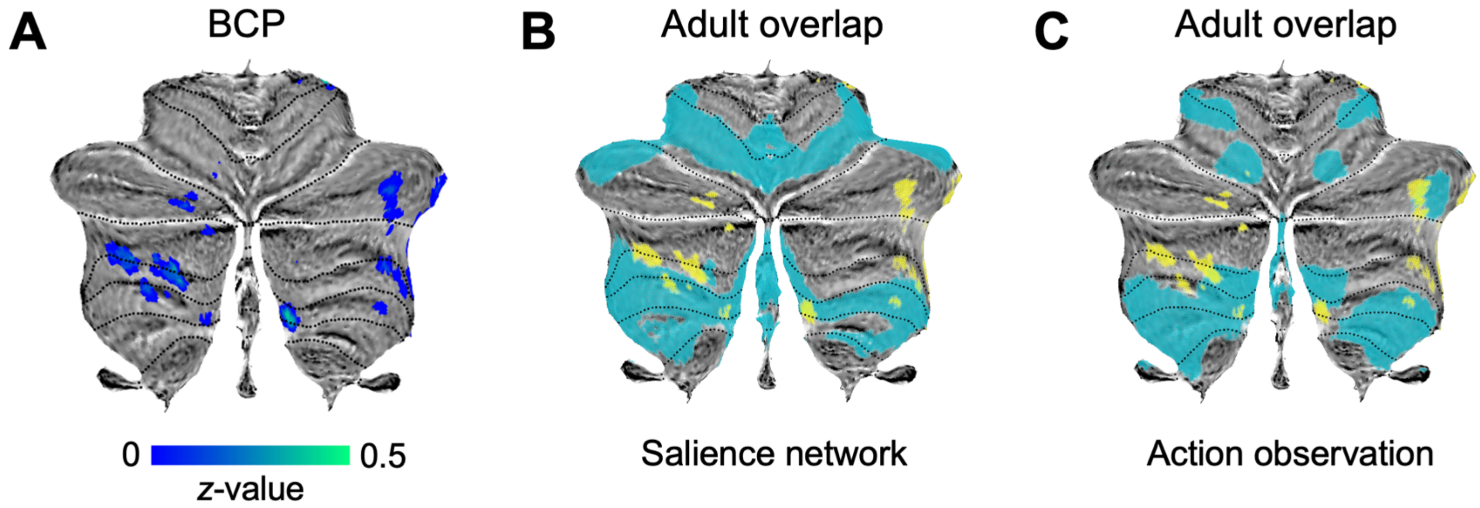


**Supplementary Figure 4.** Structural correlates of joint attention in the developing cerebellum. **A.** Grey matter changes associated with joint attention in the Baby Connectome Project (BCP) dataset, controlling for age, sex, and total cerebellum size, *z*-scored and FDR-corrected at $q = .05$. **B, C.** Overlap of joint attention clusters with the adult salience resting-state network[1] (**B**) and the action observation map in the multi-domain task battery (MDTB) cerebellar atlas[2] (**C**). Child and adult maps are presented in yellow and blue, respectively.

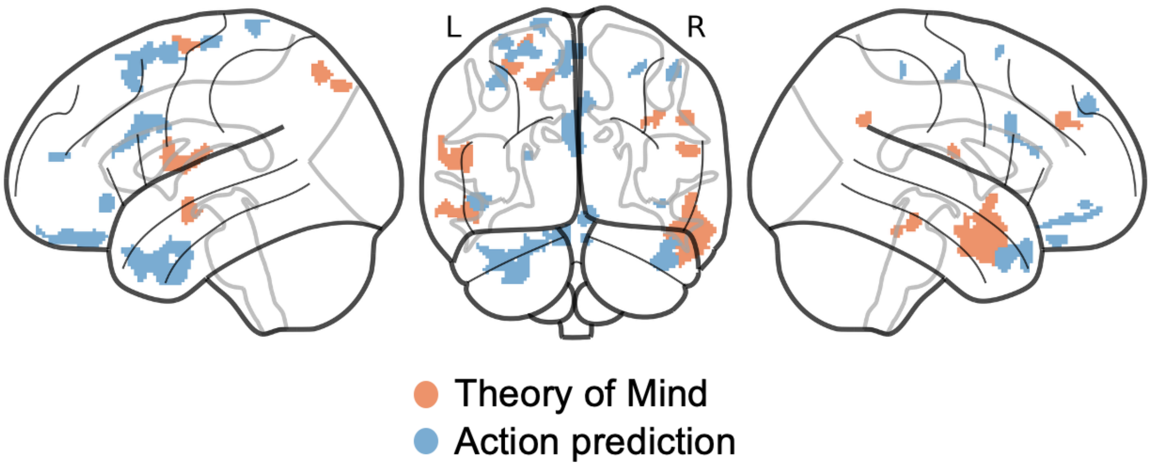


**Supplementary Figure 5.** Dissociation of explicit ToM (orange) and nonverbal action prediction (blue) cerebello-cerebral covariance clusters. Nonverbal action prediction clusters are combined for the three cerebellar ROIs (left VIIB, right VIIB, and vermis).

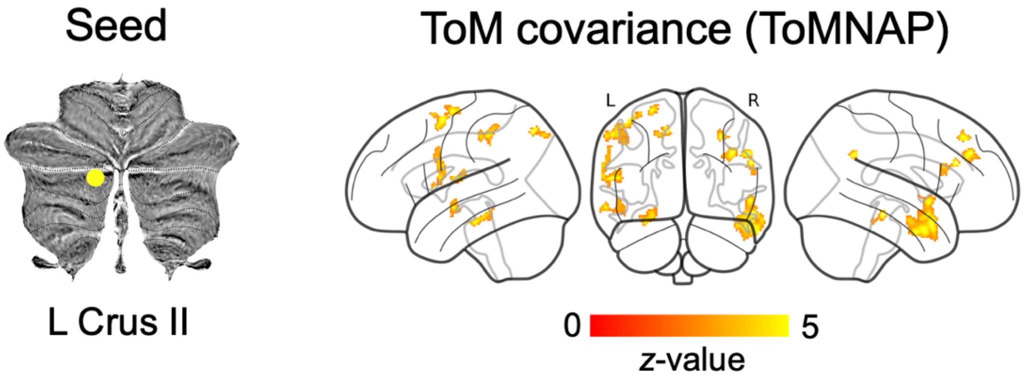


**Supplementary Figure 6.** Structural covariance between explicit ToM cerebellar ROI (left Crus II) and cerebral grey matter, controlling for age, sex, total cerebellum size, general cognitive, executive, and linguistic abilities, and nonverbal action prediction task performance in the ToMNAP dataset ($z$-scored and FDR-corrected at $q$ = .05).

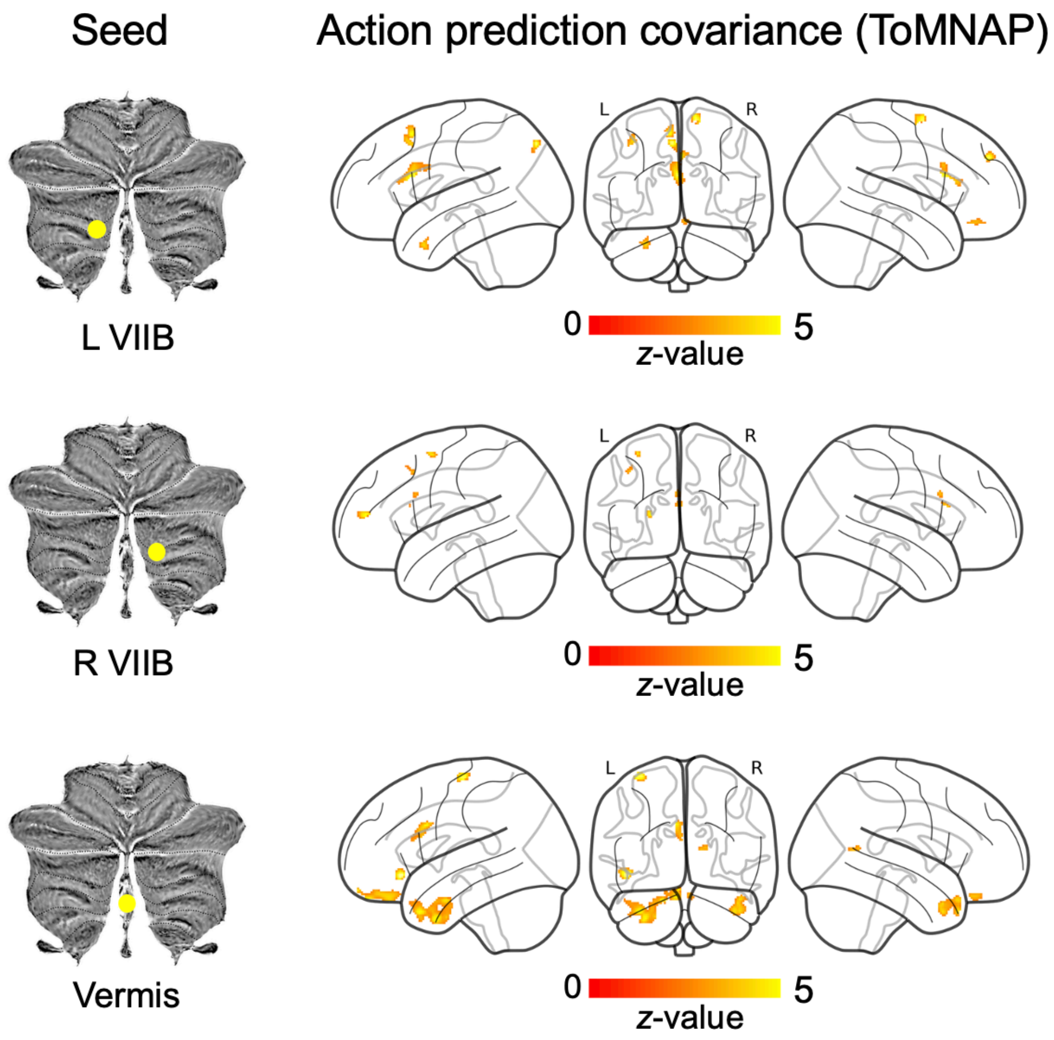


**Supplementary Figure 7.** Structural covariance between nonverbal action prediction cerebellar ROIs (left VIIB, right VIIB, and vermis) and cerebral grey matter, controlling for age, sex, total cerebellum size, general cognitive, executive, and linguistic abilities, and explicit ToM task performance in the ToMNAP dataset (*z*-scored and FDR-corrected at $q = .05$). Abbreviations: L = left; R = right.

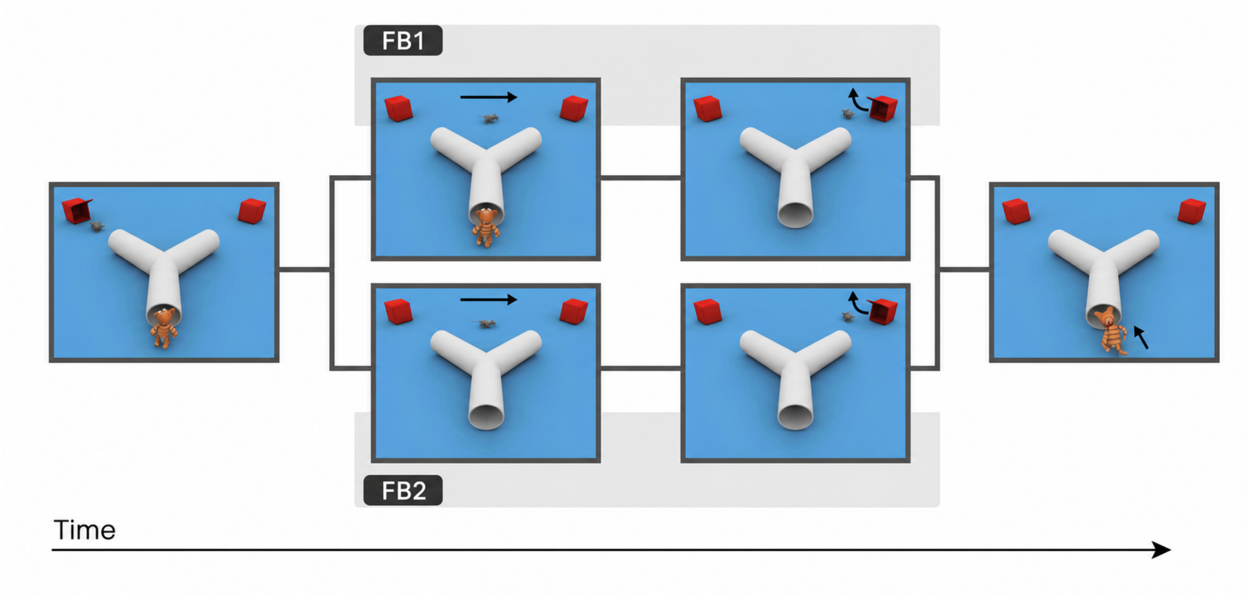


**Supplementary Figure 8.** Nonverbal action prediction task. Selected scenes from the two false belief conditions of the nonverbal action prediction task, as described in the Methods. In the present study, we focused on the condition FB1. Arrows show the movement of the animals. Adapted from Grosse Wiesmann et al.[3,4].

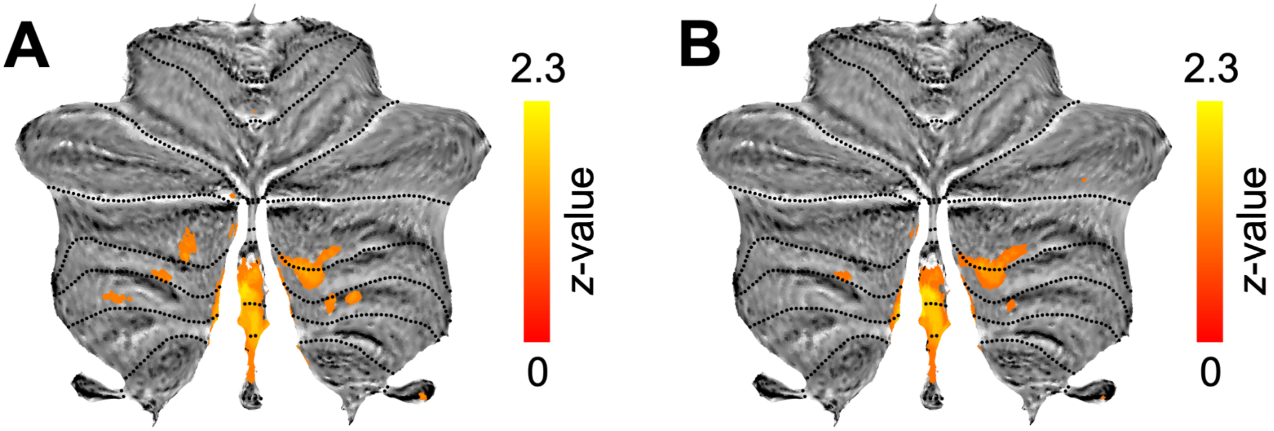


**Supplementary Figure 9.** Grey matter increases associated with nonverbal action prediction task performance (combined FB1-FB2 conditions), controlling for age, sex, total cerebellum size (**A**), and additionally general cognitive, executive, and linguistic abilities, and explicit ToM task performance (**B**) (*z*-scored and FDR-corrected at $q = .05$).

**Supplementary Table 1. Peak cerebral MNI coordinates and Neurosynth associations of ToM cerebello-cerebral covariance in the ToMNAP dataset**

| Anatomical region | Peak MNI coordinates | | | Neurosynth associations |
|---|---|---|---|---|
| | x | y | z | |
| R TPJ | 53 | -53 | 31 | Theory of Mind |
| R STSa | 59 | -1 | -30 | Theory of Mind |
| R STSb | 50 | 4 | -18 | Listening, language comprehension |
| L STS | 50 | -6 | -14 | Listening, watching, social, autobiographical |
| L operculum | 54 | -8 | 12 | Somatosensory, nociceptive |

MNI = MNI152NLin2009cAsym template; STS = superior temporal sulcus; TPJ = temporoparietal junction; R = right; L = left.

**Supplementary Table 2. Peak cerebral MNI coordinates and Neurosynth associations of ToM cerebello-cerebral covariance in the Richardson dataset**

| Anatomical region | Peak MNI coordinates | | | Neurosynth associations |
|---|---|---|---|---|
| | x | y | z | |
| L STS | -51 | -10 | -39 | Theory of Mind, perspective-taking, empathy |
| L PreC | -54 | -8 | 40 | Theory of Mind |
| R somatosensory | 54 | -11 | 15 | Somatosensory, nociceptive |
| L parietal cortex | -20 | -72 | 39 | Working memory, load |

MNI = MNI152NLin2009cAsym template; STS = superior temporal sulcus; PreC = precuneus; R = right; L = left.

**Supplementary Table 3. Peak cerebral MNI coordinates and Neurosynth associations of nonverbal action prediction cerebello-cerebral covariance in the ToMNAP dataset**

| Seed | Anatomical region | Peak MNI coordinates | | | Neurosynth associations |
|---|---|---|---|---|---|
| | | x | y | z | |
| L VIIB | L cingulate | -2 | 16 | 24 | Fear, pain, reinforcement |
| | L MFG | -6 | 18 | 57 | Supplementary motor, mentalizing, semantic |
| | L dlPFC | -37 | 18 | 48 | Autobiographical, emotion regulation, reappraisal |
| | R mOFC | 5 | 41 | -16 | Reward, social, mentalizing |
| | L temporal pole | -26 | 9 | -35 | Semantic memory, bias, selective attention |
| | R mPFC | 3 | 51 | 36 | Social, moral, mental states |
| R VIIB | L cingulate | -2 | 16 | 24 | Fear, pain, reinforcement |
| | L MFG | -6 | 18 | 57 | Supplementary motor, mentalizing, semantic |
| | L dlPFC | -37 | 18 | 48 | Autobiographical, emotion regulation, reappraisal |
| | R somatosensory | 30 | -36 | 56 | Action, action observation, movement |
| Vermis | L cingulate | -2 | 16 | 24 | Fear, pain, reinforcement |
| | L temporal pole | -26 | 9 | -35 | Semantic memory, bias, selective attention |
| | L mOFC | -4 | 46 | -25 | Social, strategy, selective attention, mentalizing |
| | R STS | 44 | 15 | -34 | Mental states, autobiographical, Theory of Mind |
| | L IFG | -46 | 32 | -9 | Comprehension, language, semantic |

MNI = MNI152NLin2009cAsym template; MFG = medial frontal gyrus; dlPFC = dorsolateral prefrontal cortex; mOFC = medial orbitofrontal cortex; mPFC = medial prefrontal cortex; STS = superior temporal gyrus; IFG = inferior frontal gyrus; R = right; L = left.

**Supplementary Methods**

***Data source.*** We sought additional evidence for cerebellar involvement in early social cognition using data from the Baby Connectome Project (BCP)[5]. Specifically, we analyzed T1w MRI data from 1–2-year-old children who completed a joint attention task, an early-emerging social ability widely considered a developmental precursor of later ToM[6,7]. The original study excluded children with preterm birth, birth complications, and medical, genetic, neurodevelopmental, or endocrine conditions; a detailed description of the inclusion and exclusion criteria is provided in Howell et al.[5]. For the present analyses, we included only children who completed the joint attention measure and who had full cerebellar coverage with no substantial motion or scanner-related artifacts after visual inspection of T1w scans. The final sample comprised 31 children ($M_{age}$ = 1.09 years, $SD_{age}$ = 0.23; 17 females). Written informed consent was obtained from a parent or legal guardian for all participants in the original study, and all procedures were approved by the relevant institutional ethics committees.

***Joint attention task.*** Participants completed an out-of-scanner assessment of responding to joint attention[8]. The task was designed to provide a dimensional measure of individual differences in joint attention performance during naturalistic social interaction. As part of the assessment, a 10-minute semi-structured parent–child interaction was video-recorded while parents were instructed to play with their child as they normally would. Trained coders subsequently evaluated children's responses to a series of hierarchically ordered joint attention bids. These bids involved progressively more redundant social cues, including gaze shifts, head turns, and other communicative signals. Children's responses were scored on a 0–4 scale according to the level of cue redundancy required to elicit a successful response, providing a continuous measure of joint attention ability.

***Data acquisition.*** MRI data were collected as part of the BCP using 3T Siemens Prisma scanners (Siemens, Erlangen, Germany) equipped with 32-channel head coils. Infants and young children were scanned during natural sleep without sedation. High-resolution whole-brain T1w structural images were acquired using a three-dimensional magnetization-prepared rapid gradient echo (MPRAGE) sequence with 0.8 mm isotropic resolution, a field of view of 256 × 240 × 166 mm, a matrix size of 320 × 300, and 208 sagittal slices. Imaging was performed with a TR of 2400 ms, a TE of 2.24 ms, and a flip angle of 8°. Detailed information regarding image acquisition and study procedures is available in Howell et al.[5].

***Data analysis.*** Image preprocessing, VBM, and structural covariance analyses were conducted using the same procedures described in the main manuscript. All statistical models included chronological age, biological sex, and total cerebellar volume (or total intracranial volume in covariance analyses) as covariates. Because the BCP data were collected across two imaging sites, the acquisition site was also included as an additional covariate. VBM results can be seen in **Supplementary Fig. 3**. No significant clusters were identified in the cerebello-cerebral structural covariance analyses after accounting for these factors.
